# Stable knockout and complementation of receptor expression using *in vitro* cell line derived reticulocytes for dissection of host malaria invasion requirements

**DOI:** 10.1101/495853

**Authors:** Timothy J Satchwell, Katherine E Wright, Katy L Haydn-Smith, Fernando Sánchez-Román Terán, Joseph Hawksworth, Jan Frayne, Ashley M Toye, Jake Baum

## Abstract

Invasion of and intracellular development within the red blood cell by malaria parasites requires interaction with a multitude of host proteins expressed at the surface of or inside the cell. Perhaps the biggest obstacle to dissection of specific functions of host proteins during invasion is the inability to manipulate protein expression in the genetically intractable anucleate red blood cell. Whilst genetic manipulation and subsequent *in vitro* differentiation of nucleated erythroid precursors have facilitated progress in this area, such approaches are limited by the finite proliferative capacity of primary hematopoietic stem cells, and a failure of erythroleukemic cell lines to enucleate, respectively. Here, we report that reticulocytes derived through *in vitro* differentiation of an enucleation competent immortalized erythroblast cell line (BEL-A) support both successful invasion and growth by *Plasmodium falciparum*. Using CRISPR-mediated gene knockout and lentiviral expression of open reading frames, we validate an essential role for the erythrocyte receptor basigin in *P. falciparum* invasion and, for the first time, demonstrate that this can be rescued by re-expression of the receptor or of a mutant thereof. Specifically, using sustainable edited clones derived from this line, we exclude a functional role for the cytoplasmic domain of basigin during invasion, and challenge the reported requirement of its associated receptor cyclophilin B. These data establish the use of reticulocytes derived from immortalized erythroblasts as a crucial model system to explore specific hypotheses regarding host receptor requirements and involvement in *P. falciparum* invasion.

## Introduction

Malaria, an infectious disease caused by *Plasmodium* parasites, is an enormous economic and health burden. Every year more than 200 million clinical cases and almost half a million deaths are reported, with most fatalities occurring in children under the age of five (WHO, 2017). Parasite invasion into and development within red blood cells is responsible for all pathology associated with this disease. Invasion begins with the interaction between a merozoite (the invasive parasite form) and the red blood cell (RBC) surface, which precedes penetration and intracellular vacuole formation via mechanisms that remain incompletely understood. One host protein implicated in the invasion process is basigin (BSG, CD147), a surface receptor believed to be essential for invasion via its interaction with *Plasmodium falciparum* Rh5 (Crosnier et al., 2011) though our understanding of the function the interaction plays in invasion is limited.

One of the biggest obstacles to the investigation of host protein involvement in red blood cell invasion is the intractability of this anucleate cell as a system for genetic manipulation. Elegant use of proteases, blocking antibodies and the identification and study of rare naturally occurring red blood cell phenotypes have provided valuable information regarding the requirement for individual receptors (reviewed in (Cowman et al., 2017; Salinas and Tolia, 2016; Satchwell, 2016). However, reliance upon the identification of often vanishingly rare blood donors to provide insight is inefficient and precludes hypothesis-driven investigation of host protein involvement in invasion.

The capacity to derive reticulocytes (young red blood cells) that are susceptible to invasion by malaria parasites through *in vitro* culture and differentiation of hematopoietic stem cells (CD34+ cells) isolated from peripheral blood or bone marrow has opened up myriad new possibilities to erythrocyte biologists. Such cells are phenotypically equivalent to *in vivo* derived reticulocytes and display functional equivalence to red blood cells (Giarratana et al., 2005; Griffiths et al., 2012; Timmins et al., 2011). Through lentiviral transduction of immature nucleated erythroblast precursors prior to differentiation it is now possible to generate enucleated reticulocytes with rare or novel phenotypes to study host cell protein requirements and involvement in invasion. The power of this approach was demonstrated in 2015 in a forward genetic screen employing shRNA mediated knockdown of blood group proteins in primary *in vitro* derived reticulocytes. This study identified important roles for CD55 and CD44 in *P. falciparum* invasion (Egan et al., 2015). Although informative, shRNA mediated depletion of gene expression frequently results in incomplete knockdowns that can mask all but the most obvious of invasion defects. Furthermore, the finite period in which transduced nucleated cells can be maintained in their undifferentiated state requires that for each repeated experiment a fresh transduction of new cells must be conducted.

Generation of immortalized erythroid cells able to proliferate indefinitely in an undifferentiated state whilst maintaining the capacity to undergo differentiation to generate reticulocytes has been a major goal of the erythroid biology field for decades. Early excitement surrounding the development of induced pluripotent stem cell lines (iPSCs) has been tempered by the observation of severe erythroid differentiation defects, expression of fetal globins and to date minimal capacity for enucleation (Dias et al., 2011; Focosi and Pistello, 2016; Trakarnsanga et al., 2014). The capability of orthochromatic erythroblasts, characterized by their condensed nuclei, to support malaria parasite entry (Bei et al., 2010; Tamez et al., 2009) has led to exploration of cell lines unable to complete differentiation as a model for invasion (Kanjee et al., 2017). For example, a recent study reported invasive susceptibility of semi-differentiated cells of the JK1 erythroleukemic cell line. These cells display a nucleated ‘polychromatic erythroblast-like’ morphology and despite supporting parasite invasion were not able to support further parasite development (Kanjee et al., 2017). Whilst these cells can provide insight into the requirement of receptors, such as basigin, for attachment and entry (Kanjee et al., 2017), the significant membrane complex remodelling and loss of protein (basigin and CD44 in particular) that occur prior to and during erythroblast enucleation (Bell et al., 2013; Griffiths et al., 2012; Satchwell et al., 2015) means that observations made using this model may not extrapolate well to anucleate red blood cells.

In 2017, Trakarnsanga et al. reported the generation of the first immortalized human adult erythroblast cell line – Bristol Erythroid Line Adult (BEL-A) (Trakarnsanga et al., 2017). Able to proliferate indefinitely as undifferentiated early basophilic erythroblasts, this line can be induced to undergo differentiation and enucleation, generating reticulocytes that are functionally identical to those derived from primary cell cultures. Expanding BEL-A cells can be lentivirally transduced with high efficiency, and are amenable to CRISPR-Cas9 mediated gene editing for the generation of stable clonal cell lines with knock out of individual and even multiple blood groups (Hawksworth et al., 2018). In addition, reticulocytes generated through *in vitro* differentiation of the BEL-A cell line, whether unedited or as edited sublines, derive from the same donor, eliminating the impact of donor variability and polymorphisms between experiments.

In this study, we exploit differentiation of the BEL-A cell line for the generation of enucleated reticulocytes, establishing the capacity of reticulocytes derived from an immortalized human adult erythroblast cell line to support invasion and growth of *P. falciparum*. By employing CRISPR-Cas9-mediated receptor gene knockout and lentiviral expression of open reading frames for complementation of invasion defects, we present a sustainable model system that allows us to interrogate the requirement for specific domains and associations of essential receptor complexes to the process of *P. falciparum* invasion.

## Results

In order to determine whether BEL-A derived reticulocytes represent a suitable model for the study of malaria parasite invasion, expanding BEL-A cells were induced to undergo terminal erythroid differentiation. After 15 days, enucleated reticulocytes were purified by leukofiltration under gravity and subjected to invasion assays, in which magnet purified *P. falciparum* schizonts were added to target cells. Reticulocytes derived from the BEL-A cell line in this manner were susceptible to invasion (Figure 1), with robust parasitemia equivalent to that observed in parallel using native red blood cells as noted by the appearance of rings 14 hours post parasite incubation. The ratio of the parasitemia observed in BEL-A derived reticulocytes to that in native red blood cells was 1.08 : 1 (n=3). To assess whether rings observed in the reticulocytes complete the intraerythrocytic replication cycle, cytospins were taken at 38h, 46h and 62h post invasion (Figure 1). Representative images of trophozoites, schizonts and new rings are presented, confirming the capacity of BEL-A derived reticulocytes to undergo growth and reinvasion.

**Figure 1.**
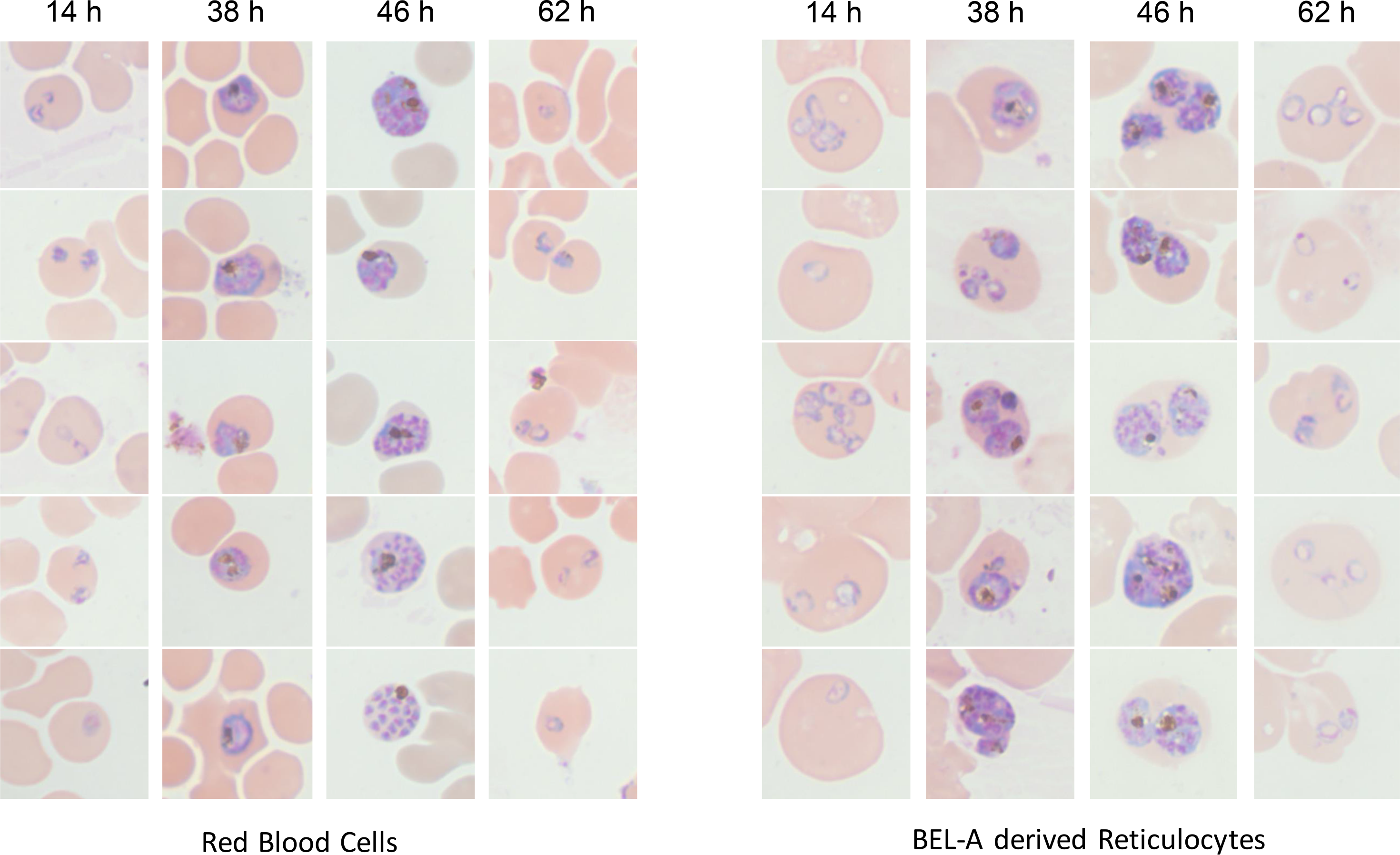
BEL-A derived reticulocytes support invasion and development of *Plasmodium falciparum*. Representative images of Giemsa stained cytospins depicting *P. falciparum* D10 ring stage parasites following successful invasion of donor erythrocytes or BEL-A derived reticulocytes, development of trophozoites, schizonts and appearance of new rings indicating reinvasion.

Since the immortalised nature of the BEL-A cell line enables use of CRISPR-Cas9 mediated gene editing for knockout of individual or multiple proteins and clonal selection, we next sought to validate use of CRISPR-Cas9 editing in the context of invasion studies. Cells were transduced with a lentiviral vector co-expressing Cas9 and a guide targeting the *BSG* gene. Transduced cells were puromycin selected and a population in excess of 80% was found to display a null BSG phenotype based on flow cytometric assessment with the monoclonal antibody HIM6. Individual null cells were FACS-sorted on this basis into 96 well plates, expanded and the null phenotype verified by flow cytometry and immunoblotting. Compound heterozygous mutations within the vicinity of the guide site were confirmed by Sanger sequencing with subsequent ICE (Influence of CRISPR Edits) software analysis (Supplemental Figure 1) (Hsiau T, 2018). BEL-A cells expanded from the selected clone were differentiated to verify capacity for enucleation; complete absence of basigin on reticulocytes was confirmed by flow cytometry (Figure 2A), whilst expression of other known malaria receptors GPA, GPC, band 3, CD55 and CD44 was unaffected as compared to unedited BEL-A derived reticulocytes (Figure 2B).

**Figure 2.**
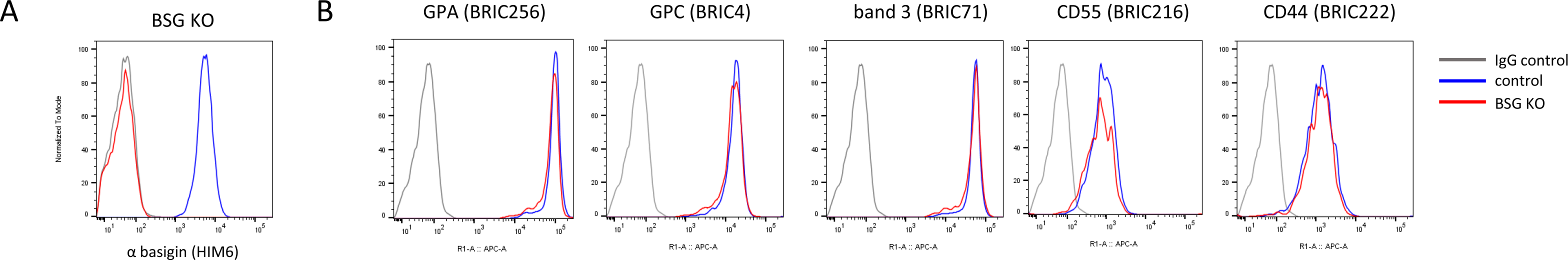
CRISPR-mediated gene editing of BEL-A cells enables generation of BSG knockout reticulocytes. **A)** Flow cytometry histogram illustrates absence of basigin (HIM6) labelling in reticulocytes derived from unedited (blue) and basigin knockout (red) BEL-A cell lines compared to IgG isotype control (grey). **B)** Flow cytometry histograms illustrate unaltered expression of indicated host receptors between reticulocytes derived from unedited and basigin knockout BEL-A cells.

Whilst previous use of *in vitro* derived reticulocytes has focused exclusively upon depleting expression of candidate receptors, complementation or ‘rescue’ studies for the reintroduction of modified proteins using such a system has not been reported. To assess the capacity to successfully rescue the invasion defect brought about by absence of basigin, sequences encoding full length wild type BSG and BSG in which the cytoplasmic tail was truncated (from 40 to 5 residues) were synthesized. In each case silent mutations were introduced within the gRNA target site to generate the null clone, rendering the constructs resistant to editing by the constitutively expressed Cas9 and gRNA. Sequences were cloned into the lentiviral vector pLVX-Neo for expression in BEL-A cells.

BSG KO BEL-A cells were transduced with lentiviral vectors for expression of the wild-type (WT) or truncated basigin (BSGΔC), resulting in populations with mixed surface presentation of the BSG extracellular domain. To maximize the possibility of obtaining reticulocytes in which expression of the reintroduced BSG was equivalent to that endogenously expressed in unedited reticulocytes, transduced undifferentiated BEL-A cells were labelled with anti-basigin HIM6 and individual clones FACS sorted to match BSG expression in modified BEL-A cell lines to that of untransduced cells. In each case, selected clones were induced to undergo differentiation, with capacity for enucleation verified and expression of BSG as well as other receptors assessed. Figure 3A shows 89.3 +/− 5.2% rescue of reticulocyte surface expression upon reintroduction of WT BSG with 98.3 +/− 4.2% for BSGΔC. Complete absence of basigin expression in BSG KO reticulocytes and altered electrophoretic mobility of BSGΔC was confirmed by immunoblotting (Figure 3B). Expression of other parasite-associated erythrocyte surface receptors was not substantially altered (Figure 3C and Supplemental Figure 2).

**Figure 3.**
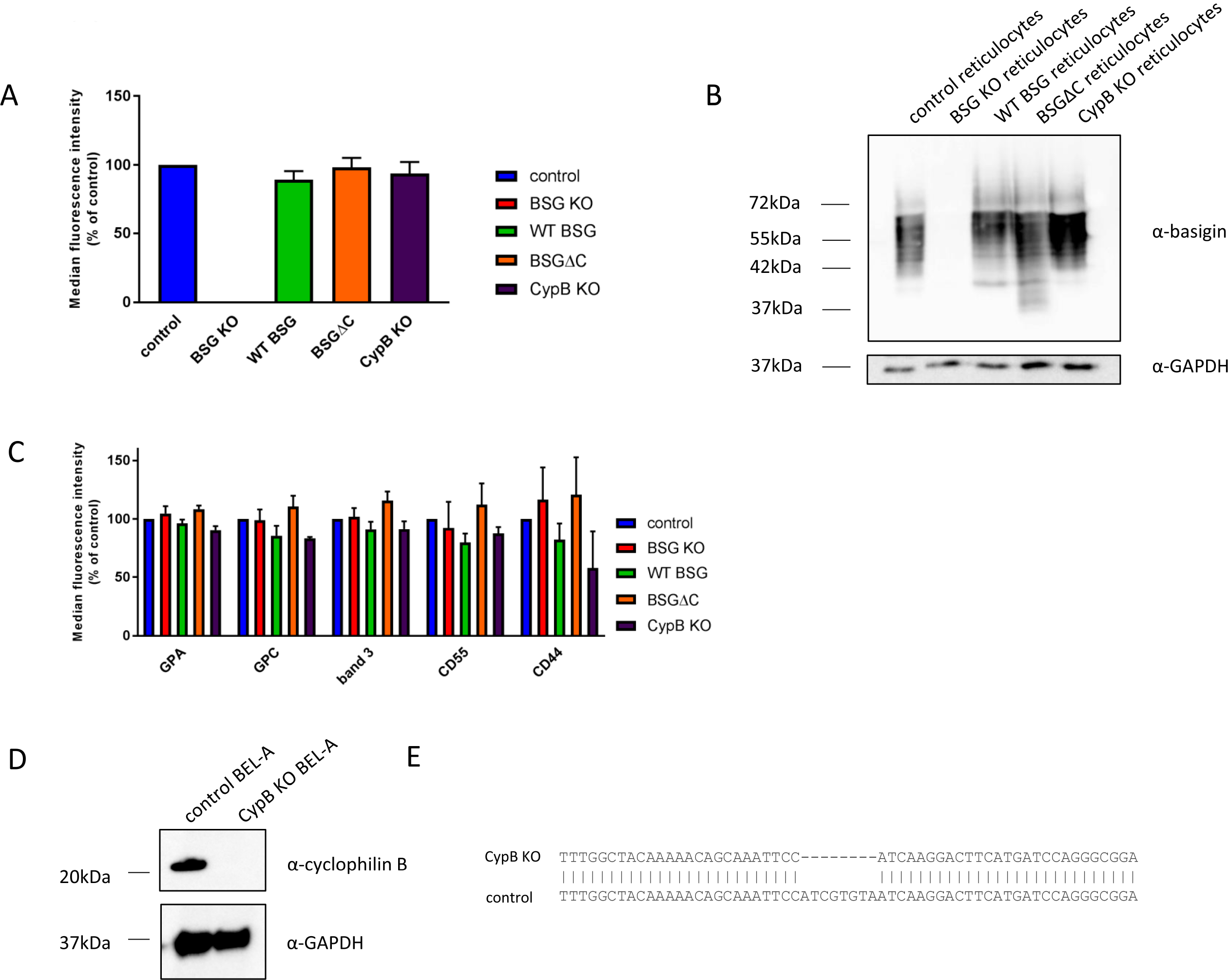
Lentiviral expression of exogenous basigin in BSG KO BEL-A cells allows for efficient rescue of receptor presentation in reticulocytes. A) Bar graphs illustrating expression of basigin as assessed by flow cytometry on reticulocytes derived from indicated cell lines. Data are normalized to endogenous expression of basigin in reticulocytes derived from unedited BEL-A cells and depict the average median fluorescent intensity across 3 independent cultures. Error bars represent standard deviation of the mean. **B)** Immunoblots of basigin and GAPDH in reticulocytes derived from indicated BEL-A cell lines. **C)** Bar graphs illustrating expression of reported malaria receptors on reticulocytes derived from indicated cell lines. Data are normalized to endogenous expression of each receptor in reticulocytes derived from unedited BEL-A cells and depict the average median fluorescent intensity across 3 independent cultures. Error bars represent standard deviation of the mean. **D)** Immunoblots of lysates from undifferentiated control (unedited) and PPIB (CypB) knockout BEL-A cells with anti-cyclophilin B and anti-GAPDH antibodies demonstrate absence of cyclophilin B expression in PPIB CRISPR gene edited cells. **E)** Sanger sequencing of PPIB gene in edited cells reveals a homozygous 8 base pair deletion at position 284 illustrated by sequence alignment that results in a frameshift.

The immunophilin protein cyclophilin B was recently reported to associate with basigin to form a host multiprotein receptor complex that may be required for invasion (Prakash et al., 2017). No null erythroid phenotypes have been reported for cyclophilin B. Therefore, to generate novel CypB KO reticulocytes and confirm its involvement in basigin-mediated parasite invasion, BEL-A were transduced with pLentiCRISPRv2 containing a guide targeting the *PPIB* (CypB) gene. Expression of CypB in reticulocytes was undetectable by immunoblotting; however, a high level of expression was observed in undifferentiated BEL-A cells (Figure 3D). Transduced cells were puromycin selected and blind sorted to derive single clones. Knockout of CypB was confirmed by immunoblotting of edited and unedited undifferentiated BEL-A cell lysates, confirming complete absence of detectable protein. Sanger sequencing identified a homozygous 8bp deletion at position 284 resulting in frameshift (Figure 3E). Reticulocytes derived from CypB KO BEL-A cells were found to express normal levels of BSG and other malaria receptors with the exception of CD44, which was variably reduced in its expression across three independent cultures (Figure 3C).

To assess invasive susceptibility of modified reticulocytes, *P. falciparum* schizonts were magnetically purified and added to 5×10^5^ target cells in a 96 well plate. After 16 hours, cytospins were prepared, Giemsa-stained and invasion quantified by manual counting of rings. Figure 4A (left) illustrates the anticipated complete absence of invasion in BSG KO reticulocytes in agreement with previous studies (Crosnier et al., 2011; Kanjee et al., 2017). An invasion assay in which unmodified and BSG KO BEL-A derived reticulocytes were incubated with excess schizonts resulted in approximately 60% parasitemia in unedited cells, with no rings observed in BSG KO reticulocytes, confirming the phenotype even under extreme invasive pressure.

**Figure 4.**
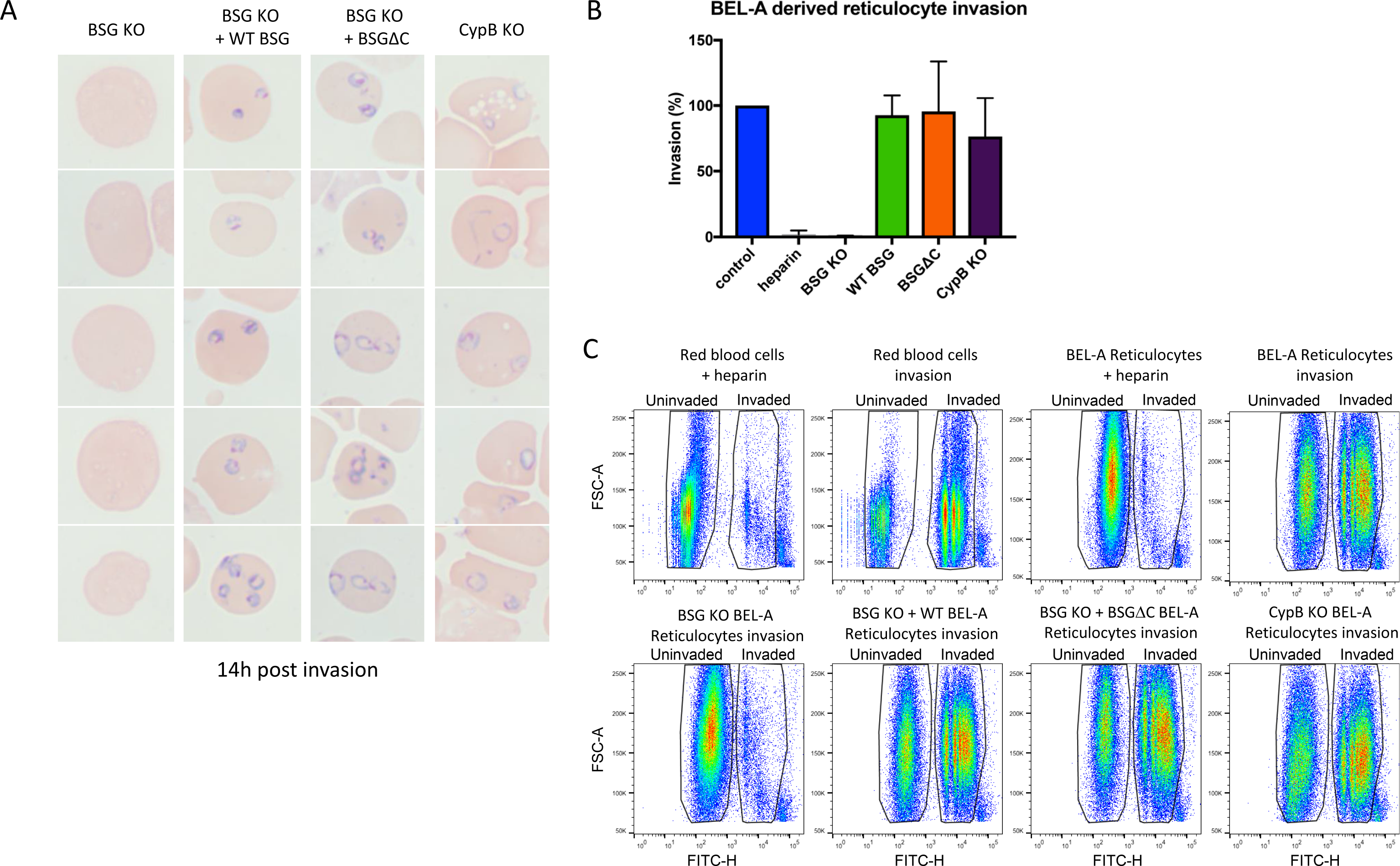
Basigin-dependent *Plasmodium falciparum* invasion of reticulocytes can be ablated and complemented through genetic manipulation of BEL-A cells. **A)** Rings are completely absent in reticulocytes derived from BSG KO BEL-A cells but are observed in reticulocytes derived from WT BSG and BSGDC rescue lines and from a CypB KO. **B)** Bar graph illustrating quantification of invasion of reticulocytes derived from indicated lines. Data represent invasion efficiency normalized to invasion in unedited control BEL-A derived reticulocytes and assessed through blinded manual counting of Giemsa stained cytospins from 3 independent experiments. Error bars represent standard error of the mean and assays in which heparin is used to inhibit invasion provide a negative control. **C)** Flow cytometric evaluation of invasion efficiency in BEL-A derived reticulocytes - dot plots illustrating capacity to identify parasitized reticulocytes following SYBR green staining.

Reintroduction of WT BSG on an endogenous BSG KO background results in the complete restoration of invasive susceptibility (Figure 4A-B). This demonstrates for the first time the capacity for complementation of invasion defects through genetic manipulation of *in vitro* derived reticulocytes. Expression of C-terminally truncated basigin rescued invasive susceptibility to that of unmodified BELA derived reticulocytes. Notably, no significant difference between unmodified reticulocytes and reticulocytes derived from the PPIB (CypB) knockout line was observed.

The ability to measure parasitemia and calculate invasion efficiency using dyes that stain nucleic acids combined with flow cytometry provides a rapid and high throughput alternative to manual counting of Giemsa stained smears or cytospins. Since this approach is reliant upon the absence of DNA in uninfected red blood cells, use of nucleated erythroid cells as a model for invasion precludes its use. *In vitro* differentiation of primary CD34^+^ cell derived erythroblasts or BEL-A cells generate enucleated reticulocytes that can be purified by leukofiltration which removes nucleated precursors and extruded pyrenocytes (Griffiths et al., 2012). To determine whether flow cytometry could be used to measure invasion efficiency in BEL-A derived reticulocytes, target cells incubated with schizonts for 16h were stained with SYBR green for 20 minutes and analysed by flow cytometry. Whilst background staining of reticulocytes is greater than that of erythrocytes (attributable to SYBR green staining of reticulocyte RNA), invaded reticulocytes could be distinguished, and invasion efficiency quantified using this approach (Figure 4C). Quantification of invasion efficiency into modified reticulocytes replicated results obtained through manual counting (Figure 4B). Of note, we observed a residual level of invasion in BSG KO cells challenged with excess schizonts that did not correlate with the complete absence of rings observed through manual counting. Manual inspection of cytospins revealed the presence of a small proportion of post-leukofiltration contaminating nucleated orthochromatic erythroblasts, together with unruptured schizonts, in quantities that correlate with the percentage of SYBR green positive events in BSG KO invasion assays. Thus, whilst this approach is best applied to assays with high parasitemia (where the influence of contaminants is minimized) and in combination with manual inspection of cytospins for confirmation of phenotypes where identified, we demonstrate that flow cytometric quantification of invasion efficiency can be employed for rapid initial assessment of invasion phenotypes in BEL-A derived reticulocytes.

## Discussion

The ability to disrupt and functionally complement phenotypes through genetic knockout and exogenous gene expression is a cornerstone approach in genetics and cell biology however direct application of these techniques to red blood cells is precluded by their anucleate nature. The development of systems for the *in vitro* culture of nucleated erythroid precursors that can be manipulated prior to their differentiation to enucleated reticulocytes, however, has paved the way to their application within the field of red blood cell biology.

Using enucleated cells derived through *in vitro* differentiation of the recently described immortalized adult erythroblast cell line, BEL-A, we demonstrate the capacity of these reticulocytes to support invasion by and growth of the malaria parasite *Plasmodium falciparum*. Using CRISPR-mediated knockout of the gene encoding the essential host receptor basigin in BEL-As we report the generation of basigin null reticulocytes. These reticulocytes are completely refractory to invasion by *Plasmodium falciparum* confirming essentiality of this receptor for invasion (Crosnier et al., 2011). By lentiviral introduction of exogenous CRISPR-resistant wild type basigin on an endogenous basigin knockout background, we generate reticulocytes with close to endogenous levels of this receptor and observe complete rescue of invasive susceptibility in reticulocytes derived from this line, demonstrating for the first time the ability to genetically complement a *P. falciparum* invasion defect that results from absence of an essential red blood cell host receptor.

Whilst the role of the extracellular domain of basigin as a site for binding of the merozoite PfRh5 is firmly established, the molecular consequences of this binding event within the host cell remain controversial, with proposed consequences including the formation of an opening or pore between parasite and host, and a Ca^2+^ influx (Aniweh et al., 2017; Volz et al., 2016; Weiss et al., 2015). Transmission of extracellular receptor binding events to downstream intracellular processes by the cytoplasmic domain of integral membrane proteins is a common theme across biology. The role of the C-terminal cytoplasmic domain of basigin is poorly understood; it has been shown to play a signal inhibitory function at the T cell synapse (Ruiz et al., 2008) and reduces the sensitivity of intracellular store-operated Ca^2+^ to cGMP in hepatoma cells (Jiang et al., 2004; Jiang et al., 2001); however, there have been no studies describing its function in red blood cells. To explore the hypothesis that the cytoplasmic domain of basigin participates in merozoite invasion in response to PfRh5 binding of the extracellular domain, a C-terminally truncated mutant was expressed in basigin knockout BEL-A cells. Reticulocytes differentiated from this modified cell line were found to exhibit no significant difference in susceptibility to invasion by *P. falciparum*. This excludes a requirement for the cytoplasmic domain in parasite invasion.

In addition to enabling dissection of the molecular basis of host receptors with established involvement in invasion, the ability to generate novel reticulocyte phenotypes with complete deficiency of prospective receptors provides a model for the verification or exclusion of candidate receptors identified via less direct approaches. A recent report proposed the existence of a multiprotein complex between basigin and cyclophilin B within the erythrocyte membrane, with a synthetic cyclophilin B binding peptide shown to inhibit merozoite invasion (Prakash et al., 2017). Since there have been no reports of cyclophilin B knockout red blood cells that would enable direct assessment of this hypothesis, we generated a cyclophilin B knockout BEL-A cell line, from which reticulocytes were derived. In contrast to the complete ablation of invasion in basigin knockout reticulocytes, no significant difference in invasive susceptibility was observed in cyclophilin B knockout reticulocytes compared to unedited controls. By generation of this novel phenotype we thus conclude that the reported interaction between parasite protein Rhop3 and cyclophilin B is not essential for merozoite invasion.

The amenability of the BEL-A cell line to genetic manipulation including via gene editing, coupled with the ability of BEL-A derived reticulocytes to permit the entire red blood cell development cycle of the parasite, allows for wide-ranging manipulation of receptors and other host proteins involved in parasite invasion, development, and egress. Future studies will undoubtedly exploit advances in the application of CRISPR guide libraries for the generation of libraries of sustainable receptor knockout lines for the study of a variety of invasion associated phenotypes as well as alternative editing approaches for site specific modification of endogenously expressed host proteins. Notably, BEL-A derived reticulocytes express both the Duffy blood group protein and transferrin receptor (Hawksworth et al., 2018) and thus should be susceptible to invasion by other malaria species including *Plasmodium vivax* and *P. knowlesi*.

In summary, we present here data that establish reticulocytes derived through differentiation of the immortalized erythroblast cell line BEL-A as a new model system for the exploration of host protein involvement in malaria invasion. We provide evidence that *P. falciparum* merozoites are able to invade and undertake the complete intracellular development cycle within the reticulocytes derived from this line. Further, using CRISPR mediated gene knockout we demonstrate the capability to generate novel reticulocyte receptor knockout phenotypes, recapitulating known invasion defects and challenging indirect evidence in support of others. Through lentiviral expression of wild type and truncated basigin on a background of endogenous protein knockout, we also present the first demonstration of complementation of a receptor-associated invasion defect whilst excluding a role for the cytoplasmic domain of basigin during this process. Overall, this establishes a model system that will enable detailed dissection of host protein involvement in multiple aspects of malaria parasite pathology.

## Supporting information

## Acknowledgements

This research was funded by a National Institute for Health (NIHR) grant to support a NIHR Research Blood and Transplant Unit (NIHR BTRU) in Red Blood Cell Products at the University of Bristol in Partnership with NHSBT (NIHR-BTRU-2015-10032; TJS, JF, AMT), grant funding from NHS Blood and Transplant R&D committee (NHSBT WT15-05; TJS, KLH-S, AMT) and Wellcome (Investigator Award 100993/Z/13/Z to JB). KEW is supported through a Henry Wellcome Postdoctoral Fellowship (107366/Z/15/Z), JH was funded by a EPSRC/BBSRC SynBio Centre CDT PhD with Defence Science and Technology Laboratory as an industrial partner. The views expressed are those of the author(s) and not necessarily those of the NHS, the NIHR or the Department of Health.

## Author contributions

Experiments were conceived and designed by TJS and KEW. TJS and KEW carried out the majority of experiments, performed the analysis and prepared the figures. KLH-S, FS-RT, JH assisted with experimental work, JF provided BEL-A cell line. AMT and JB contributed equally in supervision of the project and edited the manuscript. TJS and KEW wrote the manuscript. All authors read and approved the manuscript.

## Methods

### Cloning

Lentiviral vector pLentiCRISPRv2 containing guide sequence TTCACTACCGTAGAAGACCT targeting BSG or TGAAGTCCTTGATTACACGA targeting PPIB (CypB) was ordered from Genscript. Lentiviral expression constructs for complementation experiments were generated using a pLVX-Tight-Puro plasmid modified to contain a CMV enhancer and promoter and a neomycin resistance gene. Gibson assembly was used to combine the NotI-linearized plasmid with sequence encoding either full-length BSG, or BSG lacking the cytoplasmic domain (residues 1-234; BSGΔC). The human BSG gene was codon reoptimized, including six mutations in the CRISPR guide sequence used to knock out BSG (new sequence 5′-TTTACCACCGTGGAGGATCTGG-3′). The re-optimized gene was synthesized commercially with flanking sequences (5′ flank CTAGCGCTACCGGTCGCCACCGGATCCACC; 3′ flank GCGGCCGCGCCGGCTCTAGATCGCG) for Gibson assembly to generate the full-length BSG construct. To generate the BSGΔC construct, PCR, using the full-length synthetic gene as a template, was used to add sequences for Gibson assembly (primers 5′-CTAGCGCTACCGGTCG CCACCGGATCCACCAT GGCCGCCGCCCTCTTT GTC-3′ and 5′-CGCGATCTAGAGCCGGCGCGGCCGCTCACTTCCGCCGCTTCTCGTAGATG-3′).

### BEL-A Cell Culture

BEL-A cells were expanded, differentiated and leukofiltered as described previously (Hawksworth et al., 2018)

### Lentiviral Transduction

Lentivirus was prepared according to previously published protocols (Satchwell et al., 2015). For transduction of BEL-A cells, concentrated virus was added to 2 × 10^5^ cells in 2 ml medium in the presence of 8 μg/ml polybrene for 24 h. Cells were washed three times in PBS and resuspended in fresh medium. For vlentiCRISPR v2 transductions cells were selected 24 hours after removal of virus using 1 μg/ml puromycin for 48 h.

### Selection of Individual Clones by FACS

BEL-A cells transduced with vlentiCRISPRv2 with guide targeting BSG were immunolabelled with propidium iodide and anti-basigin antibody HIM6. Individual cells within the negative population FACS were sorted into a 96 well plate for onward culture using a BD Influx Cell Sorter. To derive clones of BSG knockout cells transduced with pLVX constructs in which basigin expression was matched to endogenous levels, transduced populations were immunolabelled with HIM6 and single clones FACS isolated through matching to a tight gate based on endogenous basigin expression of unedited BEL-A cells. Derivation of PPIB (CypB) knockout clones was achieved through blind sorting of individual clones followed by downstream screening using Sanger sequencing and immunoblotting.

### Flow Cytometry

For flow cytometry on undifferentiated BEL-As, 1 × 10^5^ cells resuspended in PBSAG (PBS + 1 mg/ml BSA, 2 mg/ml glucose) + 1% BSA were labelled with primary antibody for 30 min at 4°C. Cells were washed in PBSAG, incubated for 30 min at 4°C with appropriate APC-conjugated secondary antibody, and washed and data acquired on a MacsQuant VYB Analyser using a plate reader. For differentiated BEL-As, cells were stained with 5 μg/ml Hoechst 33342 then fixed if required in 1% paraformaldehyde, 0.0075% glutaraldehyde to reduce antibody binding-induced agglutination before labelling with antibodies as described. Reticulocytes were identified by gating upon Hoechst-negative population.

### Antibodies

Mouse monoclonal antibodies used were as follows: BRIC4 (GPC), BRIC216 (CD55), BRIC222 (CD44), BRIC71 (band 3), BRIC256 (GPA) (all IBGRL hybridoma supernatants used 1:2), HIM6 (Biolegend 1:50), K2E2 (CypB) (Santa Cruz, 1:500), GAPDH 0411 (Santa Cruz) (1:1000), IgG1 control MG1-45 (Biolegend). Secondary antibodies were APC conjugated monoclonal anti-mouse IgG1 (Biolegend) or rabbit antimouse HRP (Dako).

### Parasite culturing

*Plasmodium falciparum* parasites of the D10 strain were maintained in human erythrocytes at between 2-4% hematocrit using standard culture conditions (Trager and Jensen, 1976).

### Invasion Assays into Erythrocytes and BEL-A derived Reticulocytes

Schizont stage parasites were magnetically purified using the Magnetic Cell Separation (MACS) system (Miltenyi Biotec) (Boyle et al, 2010) and added to wells of a 96-well plate containing either erythrocytes or leukofiltered BEL-A derived reticulocytes in culture medium. Each well contained 5×10^5^ cells, with cell numbers counted using a hemocytometer. Heparin (100 mU/μl final) was used to inhibit invasion in negative controls. After approximately 16 h, invasion was quantified using cell counting and flow cytometry. For cell counting, 1.5 ×10^5^cells were applied to a slide using a cytocentrifuge. Slides were immersed in 100% methanol fixative (15 min), Giemsa stain (10 min), and water (3 min), and imaged using a Leica DMR microscope fitted with a Zeiss AxioCam HR camera.

For flow cytometry, cells were washed in PBSAG, stained with SybrGreen (1:1000 in PBS; Sigma-Aldrich) for 20 minutes at room temperature in the dark, and washed three times in PBSAG. 1 ×10^5^ cells from each well were acquired using the FITC channel of a BD Fortessa flow cytometer.

