## Supplementary material for "Stable knockout and complementation of receptor expression using *in vitro* cell line derived reticulocytes for dissection of host malaria invasion requirements"

**Supplemental Figure 1**

**CRISPR-edited BSG KO BEL-A cell line contains compound heterozygous BSG mutations. A)** Sanger sequencing of clonal edited line shows mixed spectra emerging (vertical dashed line) within the vicinity of the gRNA target site (horizontal line), indicating a heterozygous edit. **B)** ICE analysis was unable to infer any wild-type sequence, indicating a compound heterozygous edit. Regression scores indicate likelihood of computationally generated proposed sequences in sample. Wild-type sequence is shown in red with a regression score of 0.

**Supplemental Figure 2**

**BSG KO does not reduce reticulocyte expression of other parasite associated erythrocyte surface receptors.** Flow cytometry histograms illustrating expression of indicated malaria receptors in reticulocytes derived from BSG KO, WT BSG, BSGΔC and CypB KO BEL-A cell lines compared to unedited BEL-A derived reticulocytes.

### Supplemental Figure 1

A

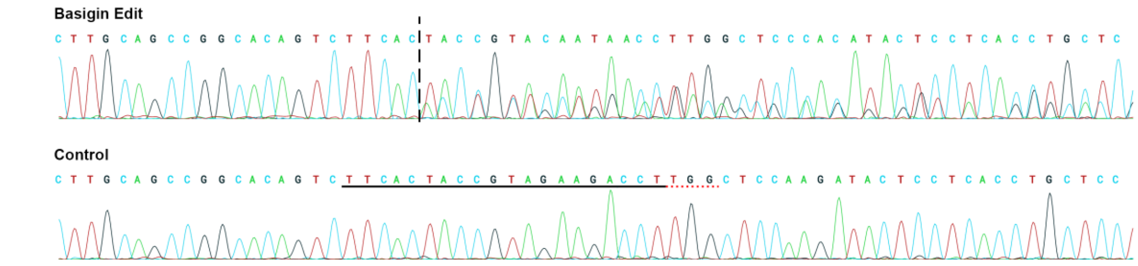


B

| | Score | ICE Proposed Sequences |
|---|---|---|
| | 0.6314 | GCACAGTCTTCACTACCGTAGAAGA \| nCCTTGGCTCCAAGATACTCCTCACCTGCTCCTTGAATGACAGCG |
| | 0.0298 | GCACAGTCTTCACT----------- \| CCTTGGCTCCAAGATACTCCTCACCTGCTCCTTGAATGACAGCGC |
| | 0.0271 | GCACAGTCTTCAC------------ \| -----------AGATACTCCTCACCTGCTCCTTGAATGACAGCGC |
| | 0.0231 | GCACAGTCTTC-------------- \| --TTGGCTCCAAGATACTCCTCACCTGCTCCTTGAATGACAGCGC |
| | 0.0216 | GCACAGTCTTCAC------------ \| -CTTGGCTCCAAGATACTCCTCACCTGCTCCTTGAATGACAGCGC |
| | 0.0166 | GCACAGTCTTCACTACCG------- \| --------CCAAGATACTCCTCACCTGCTCCTTGAATGACAGCGC |
| | 0.0119 | GCACAGTCTTCACTACCG------- \| -CTTGGCTCCAAGATACTCCTCACCTGCTCCTTGAATGACAGCGC |
| | 0.0119 | GCACAGTCTTCACTACCG------- \| CCTTGGCTCCAAGATACTCCTCACCTGCTCCTTGAATGACAGCGC |
| | 0.0108 | GCACAGTCTTCAC------------ \| ---------------ACTCCTCACCTGCTCCTTGAATGACAGCGC |
| | 0.0071 | GCACAGTCTTCACTACCGTAGAAGA \| nnnnnCCTTGGCTCCAAGATACTCCTCACCTGCTCCTTGAATGAC |
| | 0.0069 | GCACAGTCTTCAC------------ \| ----GGCTCCAAGATACTCCTCACCTGCTCCTTGAATGACAGCGC |
| | 0.0069 | GCACAGTCTTC-------------- \| ---------------ACTCCTCACCTGCTCCTTGAATGACAGCGC |
| | 0.0063 | GCACAGTCTTCACTAC--------- \| ---------------ACTCCTCACCTGCTCCTTGAATGACAGCGC |
| | 0.0062 | GCACAGTCTTCAC------------ \| CCTTGGCTCCAAGATACTCCTCACCTGCTCCTTGAATGACAGCGC |
| | 0.0056 | GCACAGTCTT--------------- \| ---------------ACTCCTCACCTGCTCCTTGAATGACAGCGC |
| | 0.0047 | GCACAGTCTTCACTACCGT------ \| ----GGCTCCAAGATACTCCTCACCTGCTCCTTGAATGACAGCGC |
| | 0.0046 | GCACAGTCTTCAC------------ \| --------------TACTCCTCACCTGCTCCTTGAATGACAGCGC |
| | 0.0040 | GCACAGTCTTCAC------------ \| -----GCTCCAAGATACTCCTCACCTGCTCCTTGAATGACAGCGC |
| | 0.0010 | GCACAGTCTT--------------- \| --------------TACTCCTCACCTGCTCCTTGAATGACAGCGC |
| WT | 0.0000 | GCACAGTCTTCACTACCGTAGAAGA \| CCTTGGCTCCAAGATACTCCTCACCTGCTCCTTGAATGACAGCGC |

### Supplemental Figure 2

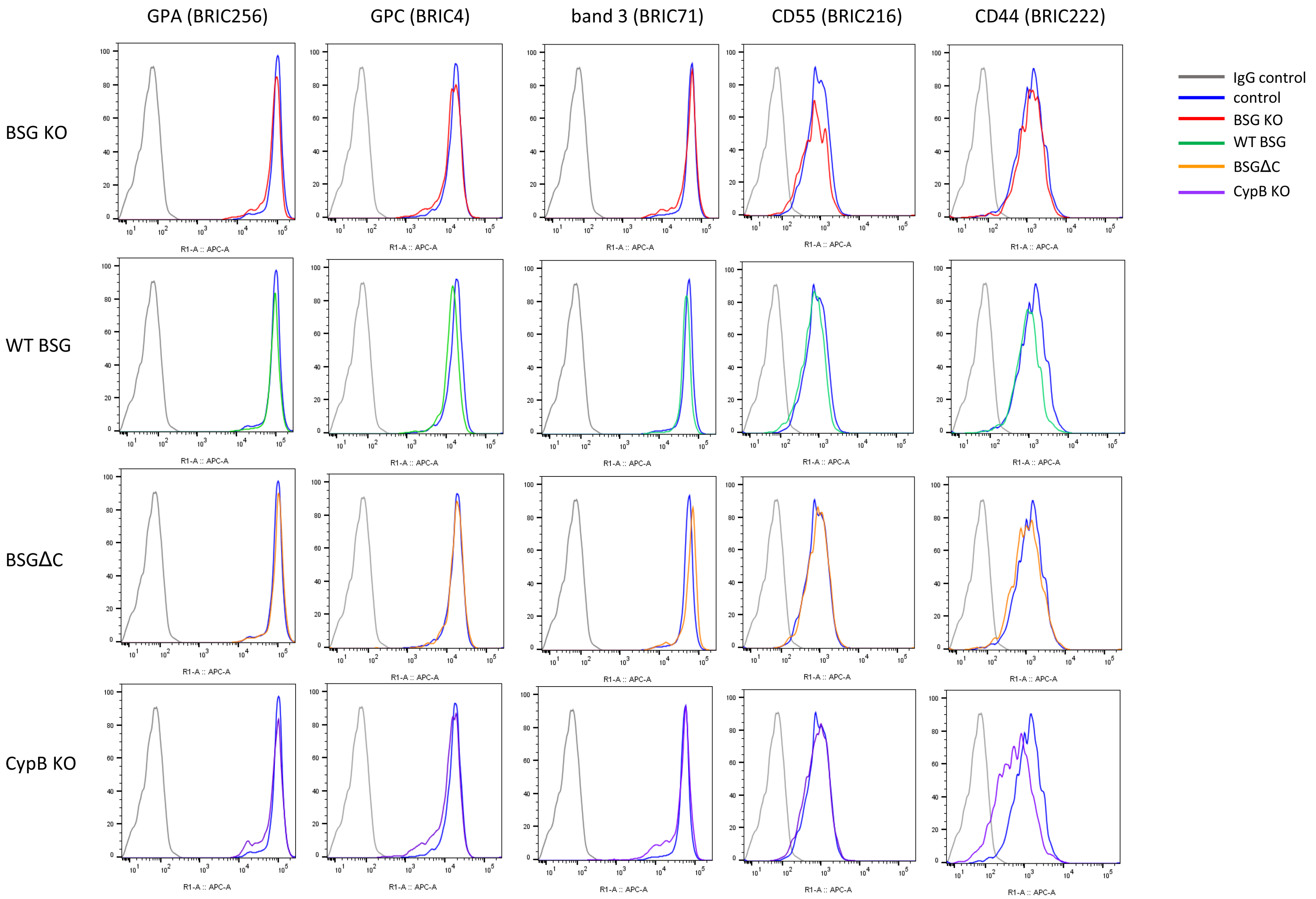
